# The spatiotemporal dynamics of bottom-up and top-down processing during at-a-glance reading

**DOI:** 10.1101/2024.02.26.582140

**Authors:** Nigel Flower, Liina Pylkkänen

**Affiliations:** Department of Linguistics, New York University; Department of Psychology, New York University

**Keywords:** top-down processing, bottom-up processing, error detection, syntax, magnetoencephalography, rapid parallel visual presentation

## Abstract

Like all domains of cognition, language processing is affected by top-down knowledge. Classic evidence for this is missing blatant errors in the signal. In sentence comprehension, one instance of this is failing to notice word order errors, such as transposed words in the middle of a sentence: *you that read wrong* (Mirault et al., 2018). Our brains seem to fix such errors, since they are incompatible with our grammatical knowledge. But how do our brains do this? Following behavioral work on inner transpositions, we flashed four-word sentences for 300ms using rapid parallel visual presentation (RPVP, Snell and Grainger, 2017). We compared their magnetoencephalography responses to fully grammatical and reversed sentences. Left lateral language cortex robustly distinguished grammatical and reversed sentences starting at 213ms. Thus, the influence of grammatical knowledge begun rapidly after visual word form recognition (Tarkiainen et al., 1999). At the earliest stage of this neural “sentence superiority effect,” inner transpositions patterned between grammatical and reversed sentences, showing evidence that the brain initially “noticed” the error. However, a hundred millisecond later, the inner transpositions became indistinguishable from the grammatical sentences, suggesting that at this point, the brain had “fixed” the error. These results show that after a single glance at a sentence, syntax impacts our neural activity almost as quickly as higher-level object recognition is assumed to take place (Cichy et al., 2014). The earliest stage involves a detailed comparison between the bottom-up input and grammatical knowledge, while shortly afterwards, knowledge can override an error in the stimulus.

## INTRODUCTION

Language comprehension involves both detailed, bottom-up analysis of a stimulus and top-down grammatical constraints (Gibson, 2006; Matar et al., 2021). The “Sentence Superiority Effect,” or SSE, is an instance of top-down syntactic knowledge guiding the interpretation of a linguistic stimulus. In studies involving the presentation of four-word stimuli at 200ms using the rapid parallel visual presentation paradigm (RPVP), participants show facilitated processing of grammatical sentences relative to scrambled sentences (Snell and Grainger, 2017; Massol et al., 2021). When the stimuli obey the grammatical rules of the participants’ native language, this syntactic knowledge can be deployed to rapidly form a sentential representation of the stimulus, requiring no longer than 200ms of presentation. This top-down knowledge also fixes stimuli containing minor word-order errors such as *you that read wrong* in studies using RPVP (Mirault et al., 2018; Snell and Grainger, 2019; Pegado and Grainger, 2020). Given that the grammatical sentence *you read that wrong* is much more likely given the sequence of words, top-down knowledge can correct the transposition of the second and third words in the ungrammatical sequence. This is evidenced by participants’ failure to notice inner transpositions in an experimental setting (Mirault et al., 2018), and the phenomenon is known as the transposed-word effect (TWE).

The literature employing the RPVP paradigm largely consists of behavioral and electroencephalographic (EEG) studies (Wen et al., 2019; Wen et al., 2021). In this study, we make use of the superior spatial resolution of magnetoencephalography (MEG) to localize the neural source of the SSE. A key objective of this study is to gain insight into how the brain “fixes” problematic stimuli containing an inner transposition. Is the bottom-up analysis of the input guided by top-down syntactic predictions in even the earliest stages, or does top-down grammatical knowledge operate slightly later in processing? Finally, we aim to offer insight into whether at-a-glance reading of multiple words involves serial (Reichle et al., 1998; Reichle et al., 2009) or parallel (Snell et al., 2018) mechanisms.

Twenty-four native speakers of English participated in an MEG experiment involving RPVP presentation of linguistic stimuli. We presented subjects with grammatical sentences, sentences with inner transpositions, and reversed sentences (see Fig 1.) to test which cortical regions show activity consistent with a neural SSE and at what point do these regions “correct” the inner transpositions and treat them as their grammatical counterparts. To query whether the brain uses serial or parallel mechanisms in at-a-glance reading of parallel input, we also used the regions discovered in the first analysis as functional regions of interest (fROIs) in a two-stage regression analysis to test whether bigram frequencies and word-to-word transition probabilities across the four-word stimulus impacted neural activity at the same time or in a more sequential fashion. We correlated bigram frequency and transition probability with the source estimates from the fROIs of each participant using simple linear models. Multiple bigram frequency effects in overlapping time windows were taken to suggest parallel processing, and multiple transition probability effects in sequential time windows serial, left-to-right processing of the multi-word stimulus (Fig. 5).

**Figure 1:**
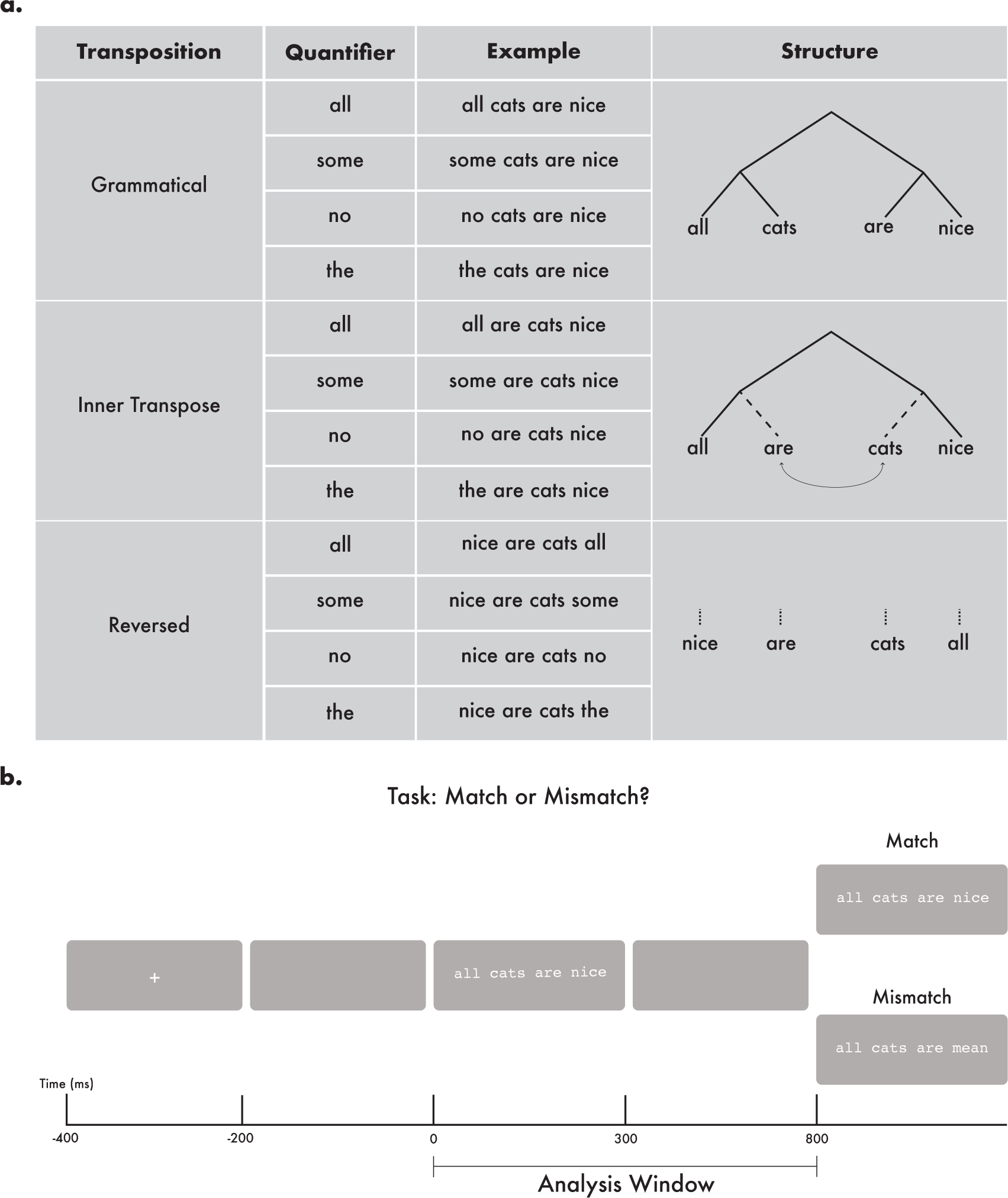
Experimental design and trial structure. (**a**) Full 3 x 4 factorial design with three sentence types (grammatical, inner-transposed and reversed) using four different types of quantifiers/determiners. Apart from adding variability to the stimuli, the quantifier manipulation was not relevant for the aims of the current study. (**b**) In our trial structure, the first, critical sentence was followed by either a matching stimulus (top) or a mismatching stimulus (bottom). prompting a match/mismatch task. MEG analysis targeted an epoch of 800ms after the onset of the first sentence.

A spatiotemporal clustering analysis revealed two clusters showing neural SSEs, the first cluster spanning over the left lateralized language cortex from 213 ms to 489 ms, and the second cluster more focalized in left inferior frontal cortex and the left anterior temporal pole from 468 – 729 ms. The early cluster showed a three-way distinction between the conditions, suggesting that top-down knowledge does not yet correct minor errors in the stimulus. However, the later cluster showed only a two-way distinction, showing a similarly increased activation for the sentences and inner transpositions as compared to the reversed sentences. This suggests that top-down knowledge of phrase structure has corrected the error in the stimulus at this later stage. Finally, our regression analysis showed that the behavior of the early cluster is consistent with parallel processing, while the behavior of the later cluster is consistent with serial processing.

## MATERIALS AND METHODS

### Participants

Thirty right-handed native speakers of English participated in the study. All participants had normal or corrected-to-normal vision and gave informed consent. Two recordings were excluded from analysis due to excessive noise, and three participants were excluded due to falling asleep. As a result, a total of 25 participants (21 women, 18-40 years old, mean age = 22.583, SD = 4.293) were included in the behavioral and MEG analyses.

### Design

To investigate the neural bases of the SSE, we employed a contrast between grammatical and ungrammatical sentences. Our Sentence Type manipulation contrasted grammatical sentences such as “all cats are nice” to reversed versions of these stimuli such as “nice are cats all.” Each of the grammatical stimuli were formed by creating a set of plural noun-adjective pairs (*cats-nice*), inserting a determiner to the left of the plural noun (*all cats-nice*) and finally by inserting the verb “are” in between the noun and the adjective. To test whether inner transpositions are detected during the composition of a parallelly presented stimulus, we also introduced another level into the Sentence Type factor for inner-transposed sentences. These stimuli were formed by taking the grammatical stimulus and simply swapping the second and third words of the sentence (*all are cats nice*). Finally, for the purposes of another project, we also varied the kind of determiner used for the sentences. A total of four determiners were chosen, *all*, *some*, *no*, and *the*, which yields the final 3 x 4 experimental design.

The nouns and adjectives in this study were selected based on character length to ensure that the sentences would be fully within the visual range of the fovea and the parafovea. Naturalness and plausibility were also considered when selecting the noun-adjective pairs for each stimulus. Each of the nouns in the study were a total of three characters long, so that they would only be four characters long in their plural form. The adjectives were either three or four characters long. The average length of the stimuli is 16.72 characters with a standard deviation of 0.84, with a min of 15 and a max of 18. The stimuli on average occupied 6.11 degrees of the visual field, meaning that most of the stimuli were close within the central visual field.

The trials began with a fixation cross on for 200ms and off screen for 200 ms. A sentence from the experimental design (see Fig 1.) was then presented for 300 ms, followed by a blank screen for 500 ms. A second sentence was then presented. This sentence was either identical to the first or involved replacing one word of the first sentence with another word taken from the lexicon used to generate the stimuli. The second sentence remained on screen until the participant marked whether the target was a match or a mismatch with respect to the first stimulus. Both the first and second stimuli were enclosed in a dark, gray rectangular box with a width of 300 pixels and a height of 50 pixels. The box was placed around the stimuli to direct participants’ gaze to the center of the screen and to help discourage eye movements outside of the boundary. The box was not presented in the intervening 500 ms between Sentence One and two. The structure of a complete trial is presented below in Figure 1.

### Procedure

Before the MEG recording, each participant had their head shape scanned with a Polhemus Fastscan three-dimensional laser digitizer to locate the positions of marker coils placed on the head during recording. The digitized head shape was used during the data preprocessing stage to constrain source localization data. Participants had the option of pausing the experiment and resting every 75 trials. The stimuli were presented to participants using the PsychoPy package in Python (Peirce, 2010) on a screen roughly 50 cm away from the participant’s face. The stimuli were presented in white Courier New font with a visual angle of 0° 34’ against a grey background. Trials were completely randomized for each participant. Participants had the option of taking a break every time they completed an eighth of the trials. Before the participants underwent the experiment, they each completed 10 practice trials to familiarize themselves with the procedure.

### MEG data acquisition and preprocessing

The raw MEG data were collected using a whole-head 157-channel axial gradiometer system (Kanazawa Institute of Technology, Tokyo, Japan) with a sampling rate of 1000 Hz. During data collection, the MEG data were filtered with a high-pass filter of 1 Hz (due to NYC environmental noise) and a low-pass filter of 200 Hz. After collection, the raw MEG data were then noise-reduced using the continuously adjusted least-square method algorithm (Adachi et al., 2001) in the MEG 160 software (Meg Laboratory 2.004A, Yokogawa Electric Corporation, Kanazawa Institute of Technology). The noise-reduced data were further low-pass filtered at 40 Hz using MNE Python (Gramfort et al., 2013, Gramfort et al., 2014). Bad channels were removed by visual inspection, and ICA was performed on the data to isolate and remove artifacts such as heartbeat and eye blinks. The cleaned data were epoched from 100 ms before the onset of Sentence One to 800 ms after the presentation of the sentence, resulting in epochs of 900 ms. The first 100 ms of the epoch was used as a baseline in the calculation of the noise-covariance matrix. Individual epochs for which any of the sensor values exceeded 3000 fT at any time point were rejected.

Estimates of source-level activity were computed from the evoked responses for each participant using dynamical statistical parameter mapping (dSPM: Dale et al., 2000). Each participant’s head shape and fiducial landmarks that were collected prior to the experiment were used to morph and co-register the “fsaverage” brain using the FreeSurfer software (http://surfer.nmr.mgh.harvard.edu/). For each condition, MEG activity was averaged, and the forward solution was computed using the Boundary Element Model (Bonnet et al., 1999, Mosher et al., 1999) as the source model. Covariance matrices were estimated using the 100 ms before the presentation of the stimulus. The inverse solution and activity at the source level was estimated by calculating minimum-norm estimates (Hämäläinen & Ilmoniemi, 1994).

### Behavioral data analyses

The behavioral data were analyzed using a 3 x 4 repeated measures ANOVA, with Sentence Type (grammatical, inner transposition, reversed) and Determiner (*all*, *some*, *no*, or *the*) as the fixed factors. For both ANOVA analyses, we first excluded any trials from the behavioral data that lasted less than 200 ms or longer than 4 seconds. We then removed any trials that deviated more than 3 standard deviations away from the grand total mean. We also chose to include reaction time and accuracy data from the participants whose MEG data were unusable, while still excluding data from the participants who were excessively sleepy. Reaction times were only analyzed for trials that were correct.

### MEG data analyses

#### Spatiotemporal Clustering Tests

Spatiotemporal cluster-based permutation tests (Maris and Oostenveld, 2007) were performed over the entire left and right cortical surfaces to detect effects of our manipulation. The p-value threshold for a cluster was set to be less than 0.05, and only clusters that spanned a minimum of 20 ms and spatial size of 10 sources were considered in the analysis. Corrected cluster p-values were estimated using 10,000 permutations. The analysis was performed over the entire 800 ms period from the onset of Sentence One till the onset of Sentence Two, to capture both early and late emerging effects.

#### Generalized Linear Model (GLM) Analysis

To probe the extent of serial vs. parallel processing in any observed effects of Sentence Type, we carried out a two-stage multiple regression model analysis by first computing a linear model of single-trial source data for each subject using the features of log bigram frequency, and log transition probability. For each subject, two models were estimated, one using a linear combination of the logged bigram frequencies of each bigram in the trial (dspm ∼ bigram(1, 2) + bigram(2, 3) + bigram(3, 4)) and one using a linear combination of the logged bigram transition probabilities of each bigram in the trial (dspm ∼ trans(1, 2) + trans(2, 3) + trans(3, 4)). Bigram frequencies and transition probabilities were computed using the Corpus of Contemporary American English (Davies, 2008).

Each of these models estimated the single-trial source data in source regions that were significant in the 3 x 4 repeated measures ANOVA of the spatiotemporal clustering analysis described above. A grand total of six models were computed for each participant. The spatiotemporal clustering analysis had identified two clusters. We thresholded the F-values of the earlier cluster to only include source points containing F-values higher than 7.5. We did the same for the later cluster but with a threshold of 5.0. The value of 7.5 was used to threshold the earlier cluster, because 7.5 was the F-value that most successfully constrained the spatial boundaries of the cluster upon visual inspection, and similarly for the later cluster with the F-value threshold of 5.0. The resulting spatial extent of the two clusters had very similar distributions to the original clusters but were much more focalized to certain regions. This was done to constrain the spatial extent of the clusters, as they both covered a wide stretch of left lateralized language cortex. The dependent measure for each of the models was the averaged dSPM values across all of the source points in the thresholded clusters.

After computing the models for each participant, we performed a temporal, one-sample t-test on the beta-coefficients of the models at each time point to see whether the coefficients were significantly different from zero. The p-value threshold for a cluster was set to be less than 0.05, and only temporal clusters that spanned a minimum of 20 ms were considered in the analysis. Cluster p-values were estimated using 10,000 permutations. We used the significant time windows from the repeated measures ANOVA analysis to constrain the search for significant effects in the temporal clustering analysis with an additional 50 ms of padding in the beginning and end of the window. For the early thresholded cluster, the time window was 163 – 519 ms, and for the late thresholded cluster, the window was 398 – 775 ms.

## RESULTS

### Behavioral results

Accuracies and RTs across sentence types and determiners are shown in Figure 2. Participants had an overall average accuracy of 89.5%. Only one participant scored below 80%, as low as 69.1%. This participant was excluded from further statistical analyses. A 3 x 4 ANOVA on the mean accuracies revealed both a main effect of Sentence Type (*F* = 8.375; *p* < 0.001) and an interaction between Sentence Type and Determiner (*F* = 3.282; *p* = 0.003). T-tests showed a significant difference between grammatical and inner transposed conditions (|*t*| = 3.802; *p* < 0.001) as well as the grammatical and reversed conditions (|*t*| = 3.33; *p* < 0.001), but not between inner transposed and reversed conditions (|*t*| = 0.476; *p* = 0.634). Further ANOVAs across each of the three sentence types were performed to find out what drove the interaction effect. When looking specifically at inner transpositions, a one level ANOVA revealed a significant effect for determiner (*F* = 6.207; *p* < 0.001), while the same analysis on grammatical sentences (*F* = 0.977; *p* = 0.403) and reversed ones (*F* = 0.77; *p* = 0.511) were both nonsignificant. This interaction was neither predicted nor relevant for the questions of the present report and is thus not further explored.

**Figure 2:**
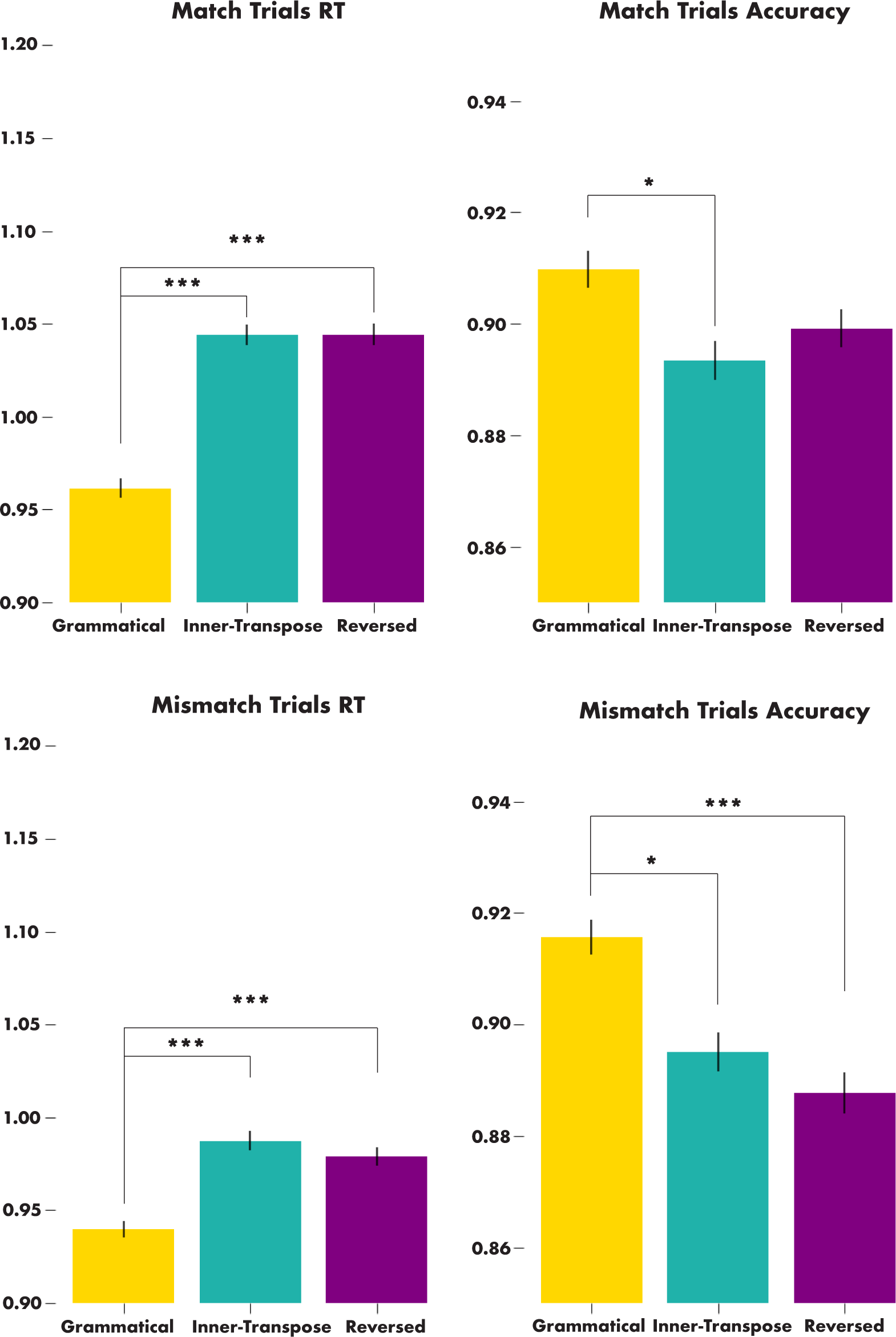
Behavioral results showing a sentence superiority effect, that is, faster and more accurate match responses to grammatical as compared to ungrammatical sentences, for both match and mismatch trials.

The mean RT for both match and mismatch trials across all participants was 1.051 seconds (*SD* = 0.585). Another 3 x 4 ANOVA was performed on RTs using the factors of Sentence Type and Determiner. The analysis revealed a significant effect of Sentence Type (*F* = 34.54; *p* < 0.001) and an interaction effect between Determiner and Sentence Type (*F* = 4.24; *p* < 0.001). A t-test showed that there was a significant difference in RT between the grammatical and inner transposition conditions (|*t*| = 7.871; *p* < 0.001) and the grammatical and reversed conditions (|*t*| = 6.5074); *p* < 0.001) but not between inner transposition and reversed conditions (|*t*| = 1.445; *p* = 0.149). As with accuracies, three more ANOVAs specific to each sentence type were carried out to investigate the interaction effect between determiner and sentence type. In line with the results from the accuracy analysis above, the one-way ANOVA looking at the effects of determiners within the sentences with inner transpositions revealed a significant effect (*F* = 7.205; *p* < 0.001), while the results for grammatical (*F* = 1.695; *p* = 0.166) and reversed sentences (*F* = 1.366; *p* = 0.251) revealed no significant effects.

### MEG results

#### Spatiotemporal clustering results

Two spatiotemporal cluster-based 3 x 4 ANOVAs were performed on source-estimated data. One was performed on the entire left cortical surface, and the other was performed on the entire right cortical surface. No significant clusters were found in the right hemisphere. In the left hemisphere, however, significant effects of sentence type were found in two separate clusters. The first cluster (Fig. 3) broadly spans over left-lateralized language cortex from 213 ms to 469 ms (p < 0.001) after the onset of the stimulus. The second cluster spans over much of the same regions as the first cluster, but with a much more focal distribution centered in the ventromedial prefrontal cortex (vmPFC) starting from 448 ms up until 725 ms (p < 0.001) after the onset of the stimulus.

**Figure 3:**
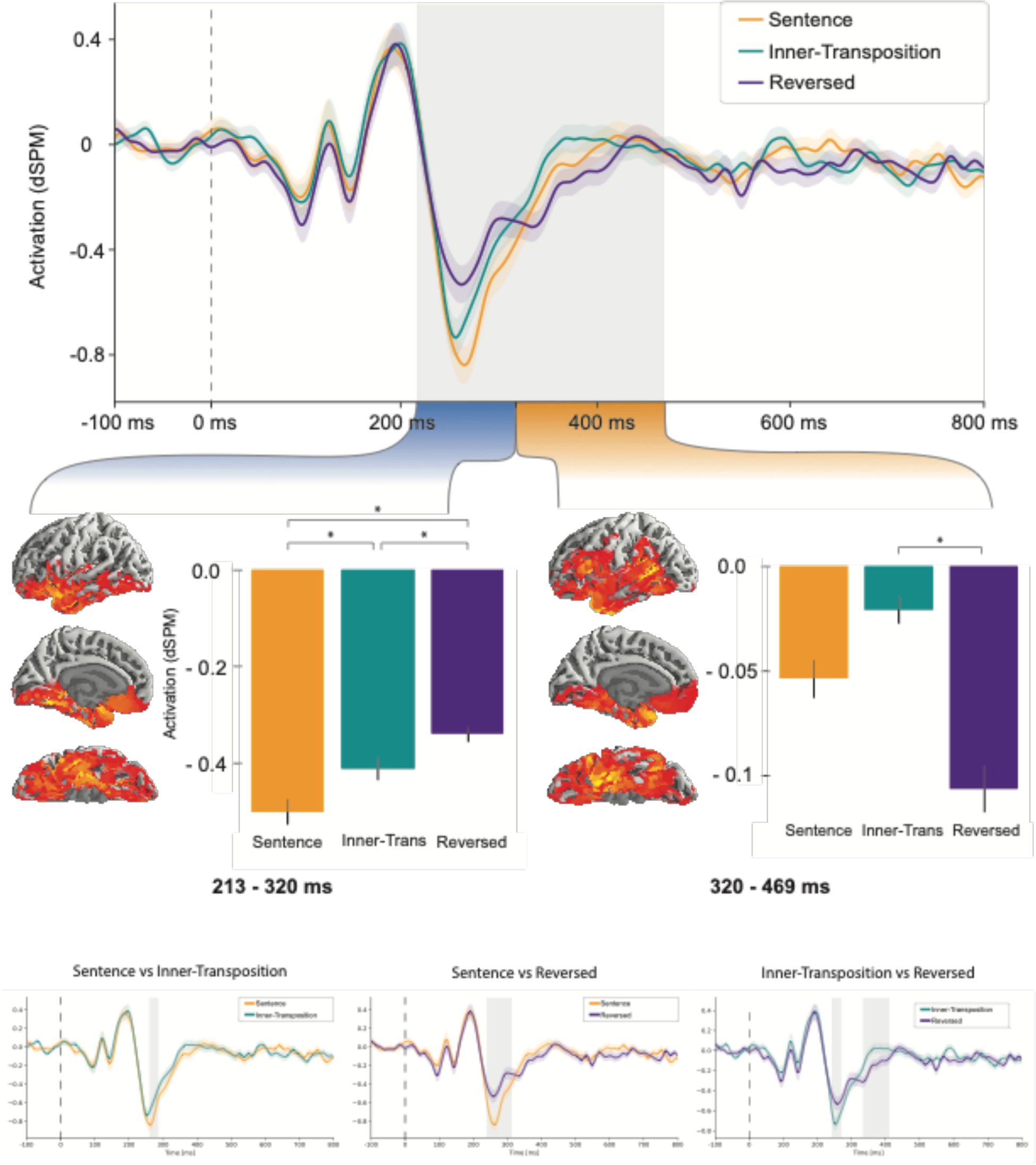
Neural sentence superiority effect in an early cluster, with an initial stage at 213-320ms showing a bottom-up profile and a later stage at 320-469ms a top-down profile.

In order to test hypotheses about the functional roles of the clusters, additional planned temporal clustering t-tests were conducted within each of them. The clusters were each used to specify fROIs in which we performed temporal clustering tests over the significant region for each cluster with extra padding of 50ms at the beginning and end of the region. The early cluster showed a three-way distinction between each of the conditions in the Sentence Type factor.

There was a significant different between the Sentence and the Inner-transpose conditions from 258 – 288ms (*p* = 0.033), a significant difference between the Sentence and the Reversed conditions from 239 – 313 ms (p < 0.001), and finally a significant difference between the Inner and Reversed conditions from 241 – 271ms (*p* = 0.0475) and from 333ms – 412ms (*p* = 0.0013). The results are consistent with a neural SSE being present in the early cluster as well as early (250 – 300ms) sensitivity to inner transpositions in the stimulus.

In the later cluster (Fig. 4), we see a different functional pattern showing only a two-way distinction. There was a significant difference between the Sentence and Reversed conditions from 508 – 533ms (*p* = 0.0491) and 536 – 605 ms (*p* = 0.0016) as well as in the Inner-transpose and Reversed conditions from 563 – 583 ms (*p* = 0.0386); however, no significant effect was found for the Sentence and Inner-transpose conditions. These results are thus consistent with the cluster exhibiting a neural SSE as well as the neural correlates of not noticing a sentence containing an inner transposition.

**Figure 4:**
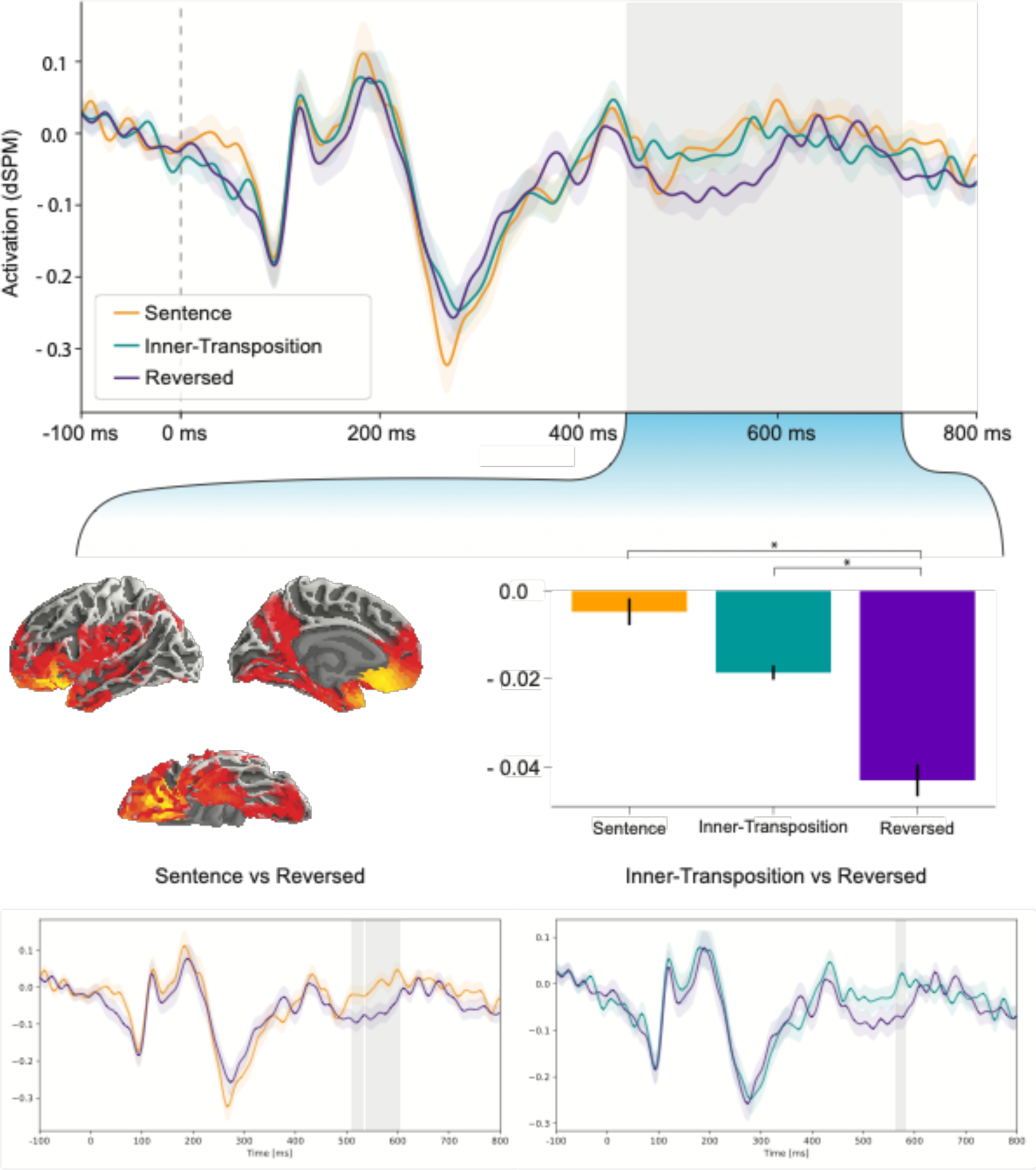
Neural sentence superiority effect in a later cluster, showing a top-down profile. Grammatical sentences and inner transpositions pattern together, both reliably diverging from reversed sentences

**Figure 5:**
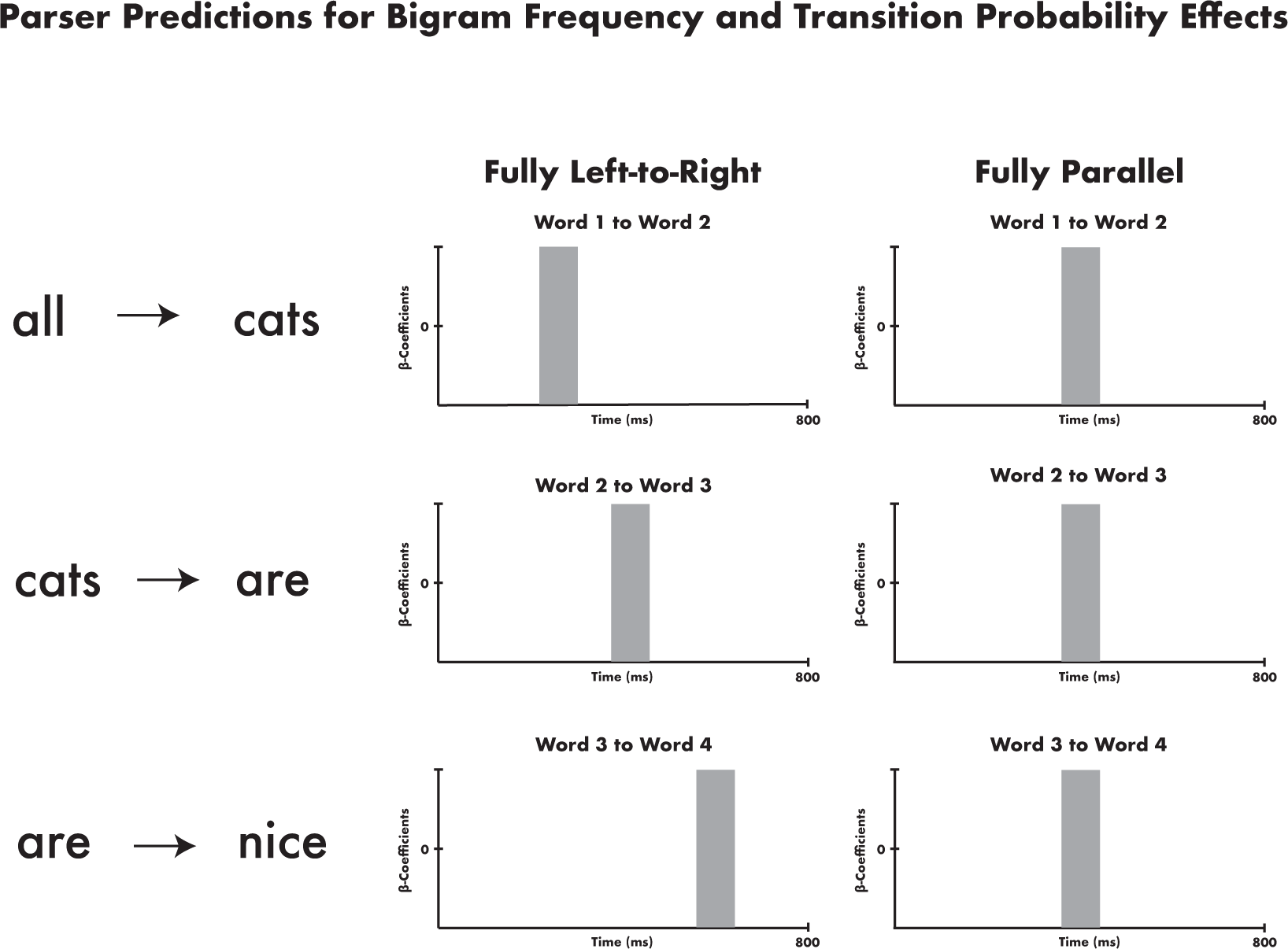
Visualization of serial and parallel hypotheses for the regression analyses of bigram frequency and transition probability reported in Figures 6 and 7.

#### Generalized Linear Model results

In order to shed light on whether the ANOVA results reported above reflect serial or parallel processing, we used the observed spatiotemporal clusters as fROI/TOIs in subsequent regression analyses testing whether bigram frequencies and word-to-word transition probabilities across the sentence affect the neural signals simultaneously or in a left-to-right sequential matter, as visualized in Figure 5. The results from the two-stage regression analyses indicate separate functional profiles for the two clusters arising from the main ANOVA analysis. The early cluster (Fig. 6) showed behavior consistent with a parallel processor, showing sensitivity to the bigram frequency of the first bigram (*p* = 0.0216, t = 238 – 274 ms) and the second bigram (*p* = 0.0181, t = 234 – 275 ms) in overlapping intervals, with the third bigram showing a trending effect shortly afterwards (*p* = 0.0659, t = 297 – 318 ms). These effects were only found in the grammatical sentences; in the inner-transposed and reversed sentences, there were no significant effects. This cluster also showed trending effects for transition probability in the grammatical sentences, but only for the second (*p* = 0.0892, t = 242 – 267 ms) and the third (p = 0.0597, t = 297 – 319 ms) bigram. As with the effects of bigram frequency, no effects were found for transition probability in the inner-transposed and reversed sentences.

**Figure 6:**
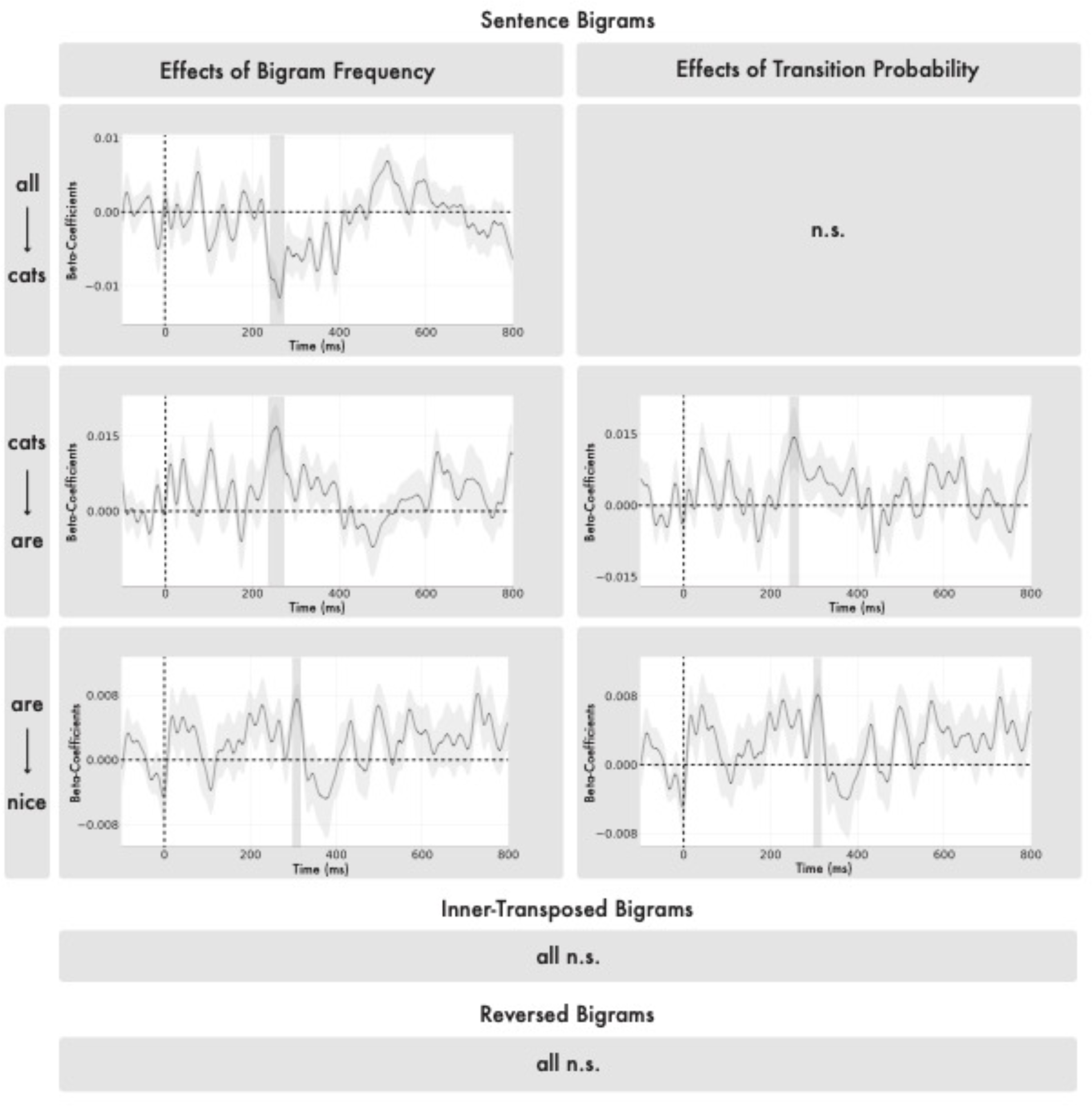
Bigram frequencies and transition probabilities affected activation in the early cluster but only for the grammatical sentences. The temporal dynamics of the significant effects fit the predictions of a parallel processor: the frequencies of the first and second bigrams were simultaneous with the third bigram immediately following. Transition probability effects were observed for the second and third word-to-word transitions and were also temporally very close.

The later cluster (Fig. 7), on the other hand, showed signs of serial left-to-right processing, but only for inner-transposed stimuli. The cluster showed effects of bigram frequency for the second bigram (*p* = 0.0118, t = 568 – 623 ms) and the third bigram (*p* = 0.0041, t = 719 – 772 ms). Similarly for transition probability, the cluster also showed effects for the second (*p* = 0.005, t = 567 – 622 ms) and third (*p* = 0.0134, t = 724 – 768 ms) bigram in roughly the same time windows. The cluster additionally exhibited an effect of transition probability for the first bigram (p = 0.0362, t = 719 – 747 ms).

**Figure 7:**
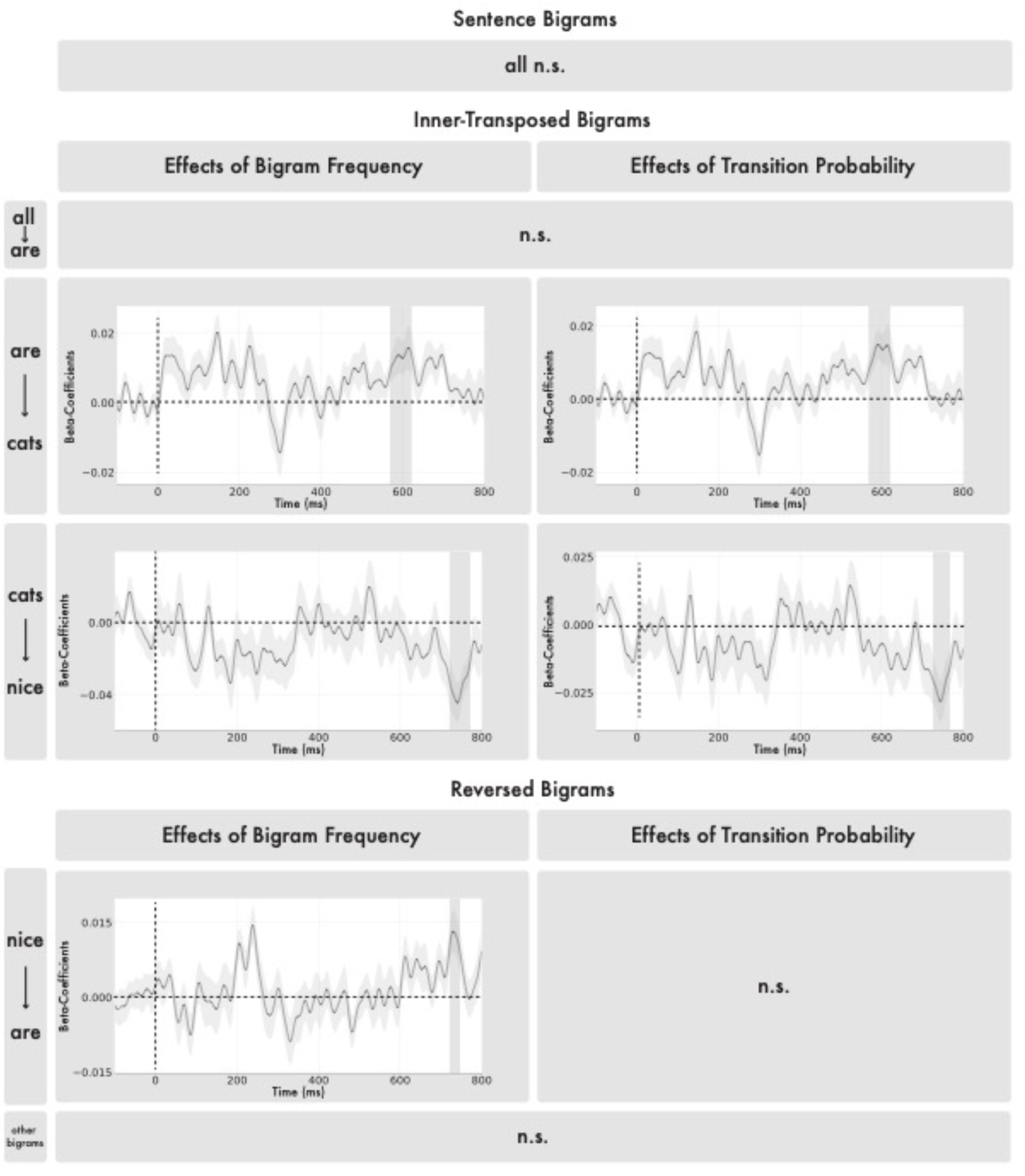
In the later cluster, bigram frequencies and transition probabilities mainly just affected the inner transpositions, where the second and third bigrams showed reliable and sequential effects of frequency of transition probability, with the third bigram following the second one by about a 100ms. Reversed sentences also showed one effect with the frequency of the first bigram having a late effect at around 750ms.

## DISCUSSION

Our perception of the world draws from both sensory-driven analysis of a stimulus as well as prior, domain knowledge of the stimulus and of the context more broadly. If a stimulus is impoverished visually, as in the case of fully gray-scaled images (Ramachandran, 1994), or auditorily, as in the case of sine-wave speech (Remez et al., 1983), then top-down knowledge is known to guide the perception of the degraded stimulus (Möttönen et al., 2006). Conversely, top-down knowledge can also “correct” our perception, causing us to misperceive or flat out ignore what would otherwise be highly salient properties of the stimulus if attention had been cued to them beforehand (Simons and Chabris, 1999; Mack and Rock, 2000). In the domain of language, a similar mechanism of top-down knowledge overwriting the literal interpretation of a stimulus is exemplified by not recognizing a transposition of an inner bigram in a sentence (Mirault et al., 2018). The underlying causes of this effect are heavily debated within the literature, with some authors arguing that the effect comes from noise during bottom-up encoding of word position (Mirault et al., 2018; Snell and Grainger, 2019), and others arguing for a post-perceptual inference mechanism fixing the transposition (Huang and Staub, 2021; Huang and Staub 2023; Hossain and White, 2023).

This study provides a neural time course of bottom-up and top-down mechanisms involved in the composition of multiword expressions by comparing grammatical sentences to sentences with the inner two words transposed and fully reversed sentences. The spatiotemporal clustering analysis revealed two separate clusters with distinct functional. The earlier of the two clusters was sensitive to both inner transpositions as well as reversals in the stimuli, suggesting that during early bottom-up processing, the brain is indeed detects phrase structure errors that may be corrected at a later stage of processing. Conversely, the later cluster showed only a two-way distinction, with the sentences and inner-transposed stimuli eliciting more activity than the reversed sentences. This suggests that during this stage of processing, the transposition has been “fixed,” because such stimuli elicited similar patterns of activation as the grammatical stimuli.

The psycholinguistic literature on the transposed-word effect posits two possible causes for the effect. In essence, the two hypotheses differ with respect to *when* the misperception of the word order arises: one positing that the bottom-up encoding of the stimulus is noisy (Mirault et al., 2018; Snell and Grainger 2019; Pegado and Grainger 2020), and the other positing that the top-down interpretation of the stimulus fixes it such that it conforms to the reader’s knowledge of phrase structure (Huang and Staub, 2021; Huang and Staub 2023; Hossain and White, 2023). The first of these hypotheses contends that word recognition takes place in parallel and that when a reader encounters a sentence such as *the white was cat big*, the noisy encoding of word position makes it possible for the reader to recognize the word *cat* before the word *was*, resulting in a TWE. Under this account (Mirault et al., 2018; Snell and Grainger 2019; Pegado and Grainger 2020), the mis-ordering arises at the level of perception, and thus the sentence would be processed bottom-up as the grammatically correct sentence *the white cat was big*.

The other account of the TWE proposes that bottom-up processing of the stimulus does encode the error, but that post-perceptual inference of the meaning of the stimulus converts it into its grammatical counterpart (Huang and Staub, 2021; Huang and Staub 2023; Hossain and White, 2023). Following Gibson et al. (2013), this approach assumes that readers use syntactic and semantic knowledge to retrieve the most likely intended message given the noisy perceived message. The findings from this study are consistent with this post-perceptual account. The behavior of the early cluster suggests that bottom-up analysis of the stimulus is sensitive to the presence of word order errors, whether they are transpositions or full reversals of the stimulus. Further, the late cluster treats the sentences and the inner-transposed stimuli identically, which suggests that at this stage or earlier, the transposition in the stimulus has been fixed such that the stimulus can be interpreted as a grammatical sentence.

Our regression analysis showed that the two clusters had complementary profiles as regards their sensitivity to bigram frequencies and transition probabilities, which we regressed against the neural data to learn about the degree of parallel versus serial processing in the early bottom-up and later top-down clusters. Although there is prior evidence that two words cannot be simultaneously recognized (White et al., 2018; White et al., 2019; White et al., 2020), the stimulus words of these studies did not linguistically compose with each other. But the sentence superiority phenomenon shows the critical impact of linguistic composition for our ability of rapid processing. Here our stimuli were clearly combinatory (grammatical sentences), clearly non-combinatory (reversed sentences) and of ambiguous nature (inner transpositions). We took overlapping effects of bigram frequencies and transition probabilities across the stimulus to be suggestive of parallel processing and sequential effects of serial processing. The early bottom-up cluster had a more parallel profile and the late top-down cluster a more serial profile. In the early cluster we observed sensitivity to the bigram frequencies of the first and second bigrams in overlapping time windows (238 – 274 ms for the first bigram and 234 – 275 ms for the second bigram) with the third bigram showing a trending effect shortly after (297 – 318 ms). These early effects were only present for grammatical stimuli and were not found for either the inner-transposed or reversed stimuli, which suggests that this early, bottom-up processing stage prefers inputs that are high in “form typicality” (Matar et al., 2021). The late cluster showed a very different profile, in which no bigram frequency or transition probability effects were elicited at all for the grammatical stimuli. Instead, this later cluster only showed effects for the inner-transposed stimuli as well as the reversed stimuli, and the temporal windows of these effects unfolded in a clearly serial fashion, unlike that of the early cluster. In sum, the regression analysis revealed a largely parallel profile limited to grammatical sentences in the early cluster and a more serial profile limited to the ungrammatical stimuli in the later cluster. These results suggest that when presented with multiple words at once, bottom-up combinatorial processing unfolds in a parallel fashion, not serially; the brain is able to make use of the parallel availability of the visual, linguistic stimulus, which is not the case in the auditory modality.

While our neural data revealed an effect of top-down linguistic knowledge on the processing of inner transpositions starting at ∼350ms after stimulus onset, at which point inner transpositions began to pattern with grammatical sentences, this was not the case in our behavioral data, where inner transpositions patterned together with the ungrammatical, reversed sentences. This is evidence of yet another stage of processing, at which the error introduced by the transposition is detected. The combination of these results and prior behavioral studies clearly suggests that the effect of inner transposition is highly task dependent, as the previous studies on Transposed-word Effects with the matching task elicited the effect by first presenting a grammatical sentence followed by the same sentence containing an inner-transposition (Mirault et al., 2018; Pegado and Grainger, 2020). However, our version of the matching task did not involve introducing a transposition on the second presentation. Our version included the possibility of participants first seeing an inner-transposed sentence to match with a second inner-transposed sentence. We made this decision to ensure a clean neural recording of a transposed sentence.

In summary, our results reveal two regions of left lateralized language cortex that show robust sentence superiority effects. At the earliest stages of multi-word comprehension, combinatorial regions of the language cortex operate on the stimulus in parallel, and later on a strictly serial type of processing takes place for ungrammatical stimuli. Furthermore, this early bottom-up processing immediately detects any deviation from grammaticality, whereas later top-down processing guided by knowledge of linguistic structure rescues stimuli that only deviate minimally from a grammatical form.

## ACKNOWLEDGEMENTS

This work was supported by award G1001 from NYUAD Institute, New York University Abu Dhabi (LP). We thank Alex White for feedback and discussion.

